# A freely accessible, adaptable hollow-fiber setup to reproduce first-order absorption: illustration with linezolid cerebrospinal fluid pharmacokinetic data

**DOI:** 10.1101/2024.12.19.629487

**Authors:** N. Prébonnaud, A. Chauzy, N. Grégoire, C. Dahyot-Fizelier, C. Adier, S. Marchand, V. Aranzana-Climent

**Author notes:** **Corresponding author:** Alexia CHAUZY, INSERM U1070 - Université de Poitiers, Pôle Biologie Santé - Bâtiment B36, 1 rue Georges Bonnet, 86022 POITIERS Cedex, France. **<u>Suggested reviewers:</u>** Ryan K Shields, Aude Ferran, Sebastian Wicha.

## Abstract

The main objective of this study was to validate an algorithm and experimental setup to simulate first-order absorption pharmacokinetic profiles without altering the standard *in vitro* hollow fiber infection model (HFIM). For that, clinical cerebrospinal fluid (CSF) linezolid concentrations after 30-minute infusions at dosing regimens 600 mg q12 h, 900 mg q12 h, and 900 mg q8 h were reproduced in the HFIM over 4 days.

To approximate the apparent first-order absorption observed on CSF pharmacokinetic profiles, we split the dosing interval into a series of sub-intervals during which continuous infusions were delivered to the system. During each sub-interval, the same amount of linezolid was delivered but the sub-intervals had different durations and flow rates which were computed by a newly developed algorithm.

In addition, we independently reproduced plasma concentrations to validate our system.

Samples were collected from the central reservoir and the extracapillary space (ECS) of the cartridge of the HFIM and assayed by liquid chromatography-tandem mass spectrometry. Observed pharmacokinetic parameters and concentrations in the ECS were compared with the target clinical pharmacokinetic parameters and concentrations.

Observed pharmacokinetic parameters were within 20 % of target pharmacokinetic parameters for all experiments, thus validating the ability of our experimental setup to reproduce plasma and CSF linezolid pharmacokinetic profiles.

The algorithm and setup are available in the open-source web application https://varacli.shinyapps.io/hollow_fiber_app/ to easily design other HFIM experiments.

## Introduction

The hollow-fiber infection model (HFIM) is a preferential model of *in vitro* pharmacokinetic/pharmacodynamic (PK/PD) study to predict bacterial killing induced by various PK profiles in order to optimize dosing regimen (1). Traditionally, PK/PD indices are determined by performing dose-fractionation studies in mouse (2). However, PK observed in animals can be different compared to that in human. For instance, drug half-life is usually faster in mouse than in human (3). Thus, HFIM is interesting because it can reproduce the PK observed in humans and over a longer duration than in *in vivo* studies which are generally limited to 24 h (1).

Most HFIM studies reproduced unbound plasma PK profiles observed after intravenous administration, and most often characterized by one or two-compartment models with first-order elimination and no absorption (4–9). Unbound plasma concentrations are considered of the best surrogate of the concentrations observed at infection sites (10). However, the antibiotic concentration at the infection site may be lower than the unbound plasma concentrations depending on the physicochemical properties of the antibiotic and to the presence of anatomical barriers (11).

This is particularly true in the context of treating cerebral infections. The brain is protected by the blood-brain barrier and the blood-cerebrospinal fluid barrier, which limit the distribution of antibiotics in the cerebrospinal fluid (CSF) (12). Although meningeal inflammation can increase the permeability of these barriers, antibiotic concentrations in CSF often remain lower and delayed when compared with those in plasma (12,13). Thus, using plasma concentrations to optimize dosing regimen can overestimate the effect at the infection site (e.g CSF) making it more relevant to reproduce concentrations observed at the infection sites in the HFIM to optimize dosing regimen.

Studies reproducing concentrations observed at infection sites (e.g CSF concentrations, lung tissue concentrations) are rare in the literature (14–16). In the studies conducted by Hope *et al*. and Kloprogge *et al*., the concentration profiles at infection sites reproduced in the HFIM were similar to those observed in plasma, which allowed them to set the programmable syringe to deliver a short infusion, as it is typically done to reproduce plasma concentrations (15,16).

However, PK profiles observed at infection sites can differ from those observed in plasma due to the presence of an ascending phase that is often slower and delayed when compared with the one observed in plasma, which cannot be mimicked by a short infusion. A way to reproduce PK profiles with absorption is to add a bottle in the HFIM to mimic a depot compartment (17). However, it consumes materials and additional medium increasing the cost of the experiment.

Moreover, the reproduction of clinical concentrations in the HFIM is not always clearly demonstrated in certain studies (14,15,18–21).

In this context, the main objective of this study was to validate an experimental setup to simulate first-order absorption PK profiles without additional compartment in an *in vitro* HFIM. Secondarily, we also wanted to validate the ability of our system to reproduce intravenous PK profiles. For this purpose, CSF and plasma concentrations of linezolid from a previous study on intensive care unit patients (ICU) (13) after three dosing regimens were reproduced in a HFIM.

## Materials and methods

A graphical summary of the methods used in this study is presented in **Figure 1**.

**Figure 1.**
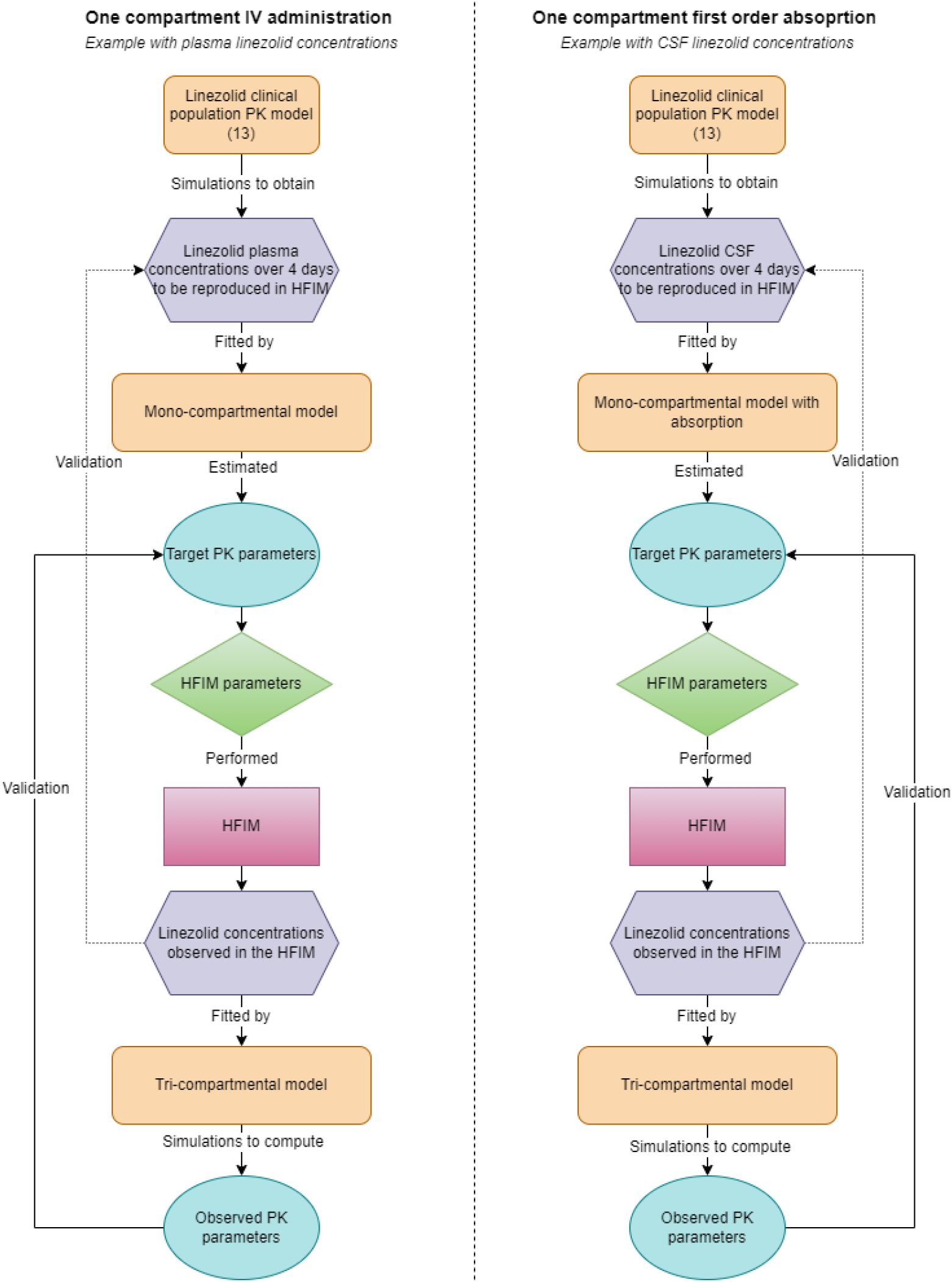
Diagram of the methods used in this study. (13) Linezolid clinical population PK model from Dahyot-Fizelier et al. Created in drawio (22).

### 1. Linezolid PK simulations

To generate typical unbound plasma and CSF linezolid concentrations in ICU patients, we used the linezolid clinical population PK model developed in a previous study (13), which aimed to evaluate the distribution of this antibiotic in the CSF of ICU patients with external ventricular drainage. Linezolid unbound plasma and CSF concentrations to be reproduced in the HFIM were simulated over 4 days for two standard dosing regimens: 30 min infusion of 600 mg every 12 h, 30 min infusion of 900 mg every 12 h, and one high dosing regimen 30 min infusion of 900 mg every 8 h. Simulations were performed using mrgsolve R package (23) with R software, version 4.2.2 (24).

### 2. Estimation of target PK parameters to reproduce plasma concentrations in HFIM

The simulated linezolid unbound plasma concentrations were fitted with a mono-compartmental model to estimate the target elimination rate constant (k_e_), maximal concentration after the first dose (C_max,1_), and concentration at steady state (C_max,ss_) using R software, version 4.2.2 (24). From the target ke value, terminal half-life (t_1/2_) and the number of doses needed to reach steady state (N_dose_) were computed. The accumulation factor (Rac) was computed from the target C_max,ss_ divided by target C_max,1_ values. Parameters are given in Error! Reference source not found..

### 3. Estimation of target PK parameters to reproduce CSF concentrations in HFIM

The simulated linezolid CSF concentrations were fitted with a mono-compartmental model with first-order absorption to estimate the absorption rate constant (k_a_), elimination rate constant (k_e_), maximal concentration after the first dose (C_max,1_) and concentration at steady state (C_max,ss_) using R software, version 4.2.2 (24). Then, the rate constants were empirically adjusted to yield a visually better match between concentrations predicted by the mono-compartmental model and concentrations from linezolid clinical population PK model (13). From the target ke value, terminal half-life (t_1/2_) and the number of doses needed to reach steady state (N_dose_) were computed. The accumulation factor (Rac) was computed from the target C_max,ss_ divided by target C_max,1_ values. Parameters are given in Error! Reference source not found..

### 4. HFIM

#### General overview of the HFIM

Linezolid unbound plasma and CSF concentrations after 30 min infusion of 600 mg every 12 h, 30 min infusion of 900 mg every 12 h and 30 min infusion of 900 mg every 8 h were reproduced in the HFIM at 37°C ± 2°C for 4 days at least in duplicate on separate occasions for each dosing regimen. A diagram of the HFIM is presented in Error! Reference source not found..

At the beginning of each experiment, stock solution of linezolid (purity > 99 % in powder form, Sigma-Aldrich, Merck KGaA, Saint-Quentin-Fallavier, France) at 10 000 μg/mL in dimethyl sulfoxide (DMSO, Merck KGaA, Saint-Quentin-Fallavier, France) was thawed from the freezer at -80°C. Then, a linezolid infusion bag was freshly prepared by diluting stock solution to the desired concentration in 0.9 % sodium chloride (Merck KGaA, Saint-Quentin-Fallavier, France). The concentration of the infusion solution depended on the dosing regimen (**Tables S1 and S2**). Then, the infusion bag was placed into an ambulatory infusion pump (Mini RythmicTM Perf+, Micrel Medical Devices S.A., Koropi, Athens, Greece) which was kept in a refrigerated box (2-10°C) throughout the experiment (**Figure 2**). The ambulatory infusion pump delivered the linezolid into a fast-flowing circulation loop connecting the central reservoir to the intracapillary space (ICS) of a dialysis cartridge (FX paed helixone dialyzer, Fresenius Medical Care, Bad Homburg, Germany) via a peristaltic pump (Masterflex L/S, Cole Parmer, Roissy, France). The flow rate in the circulation loop was set to 60 mL/min in order to obtain a rapid equilibration of linezolid concentrations between the central reservoir and the ICS of the cartridge. Extracapillary space (ECS) of the cartridge was filled with 60 mL of cation-adjusted Mueller-Hinton broth (CAMHB, Merck KGaA, Saint-Quentin-Fallavier, France) and inoculated by *Staphylococcus aureus* ATCC (American Type Culture Collection) 29213 (**Figure 2**).

**Figure 2.**
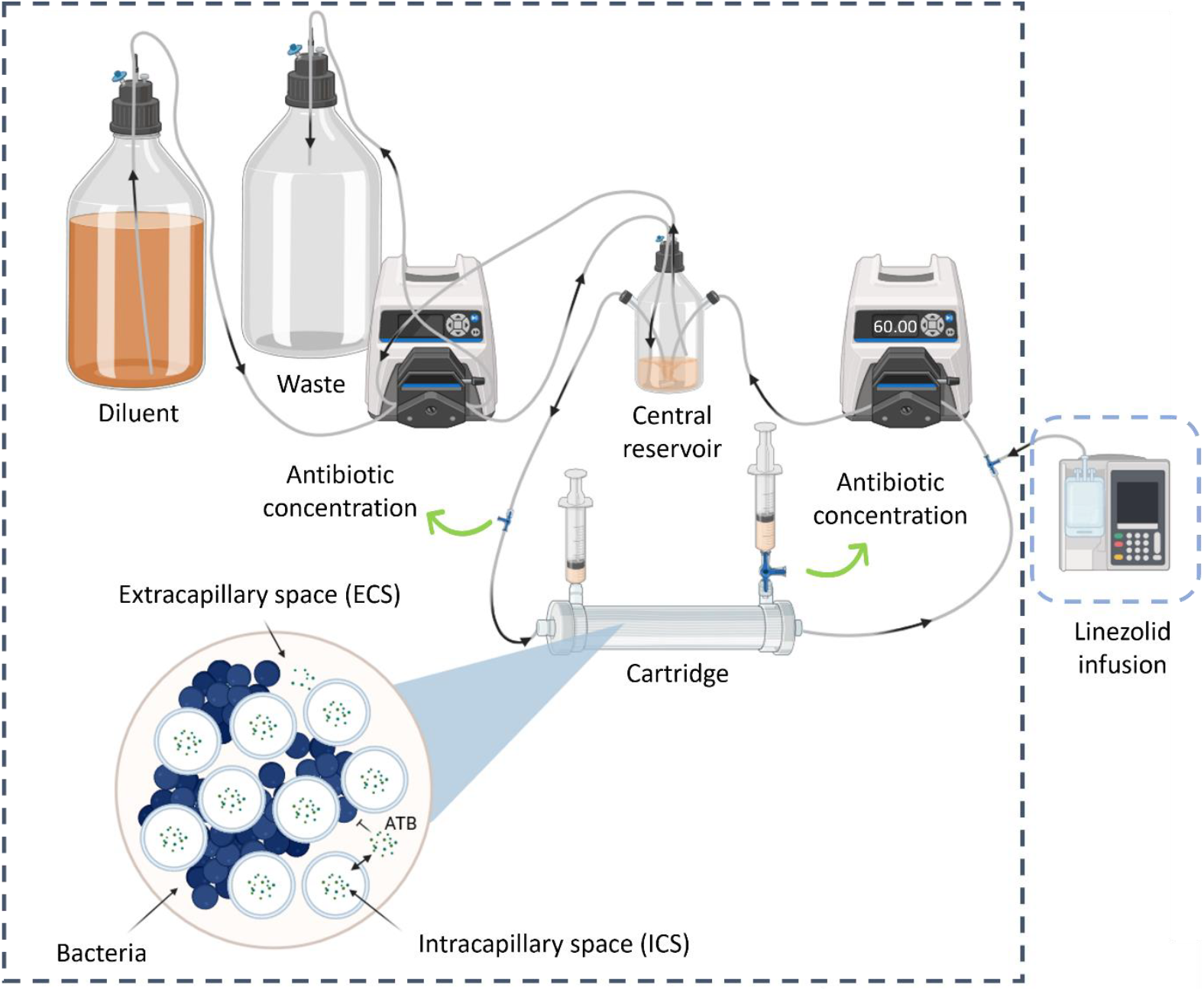
Diagram of hollow-fiber infection model. Black arrows correspond to the direction of flow. Green arrows correspond to sampling sites. Gray dashed line corresponds to the incubator at 37°C ± 2°C. Blue dashed line corresponds to the refrigerated box (2-10°C). Created in BioRender.com.

The hollow-fibers were semi-permeable polysulfone membranes with pore size varying between 1.8 nm and 3.3 nm, which allowed diffusion of linezolid between the ICS and the ECS whereas bacteria were retained in the ECS (**Figure 2**).

To mimic the elimination of linezolid, a second peristaltic pump added antibiotic free CAMHB (Diluent) to the central reservoir and removed excess CAMHB (Waste), thus diluting linezolid while maintaining a constant volume of 300 mL of CAMHB in the central reservoir (**Figure 2**)

#### Setup to reproduce linezolid plasma concentrations in HFIM

To reproduce linezolid unbound plasma concentrations in the HFIM, the ambulatory pump was set to deliver an infusion of 5 mL over 30 min every 12 h, or 8 h in the case of 900 mg q8 h. The central reservoir volume, flow rate from the central reservoir to the cartridge, cartridge volume, infusion duration and volume to be infused for one dose, dosing interval, total number of doses, experiment duration, target maximal concentration after the first dose (C_max,1_) and terminal half-life (t_1/2_) defined in **Table 1** were entered into the R-shiny application HF-App (25) to obtain the experimental parameters (flow rate from the central reservoir to the waste, diluent volume, infusion solution concentration) given in the **Table S1**.

**Table 1.** Target PK parameters for unbound plasma concentrations.

| Parameter | Dosing regimen |  |  |
| --- | --- | --- | --- |
|  | 600 mg q12 h | 900 mg q12 h | 900 mg q8 h |
| $C_{max,1}$ (mg/L) | 10.7 | 16.1 | 16.1 |
| $C_{max,ss}$ (mg/L) | 11.5 | 17.2 | 19.2 |
| Rac | 1.07 | 1.07 | 1.19 |
| $k_e$ ($h^{-1}$) | 0.230 | | |
| $t_{1/2}$ (h) | 3.01 | | |
| $N_{dose}$ | 1 | | 2 |

#### Setup to reproduce linezolid CSF concentrations in HFIM

To reproduce linezolid CSF concentrations in the HFIM, we approximated the apparent first-order absorption by a series of continuous infusions. To do so, we divided each dosing interval into 12 sub-intervals (or 8 sub-intervals in the case of 900 mg q8 h). Over all sub-intervals the same amount of linezolid was administered, only the duration and rate of administration differed between each sub-interval.

For a given target PK profile, the duration and infusion flow rates for each sub-interval was computed using the following algorithm (details about the algorithm construction are provided in **Text S1**):

1. The target maximal concentration after the first dose (C_max, 1_ in mg/L), absorption rate constant (k_a_ in h^-1^), the elimination rate constant (k_e_ in h^-1^), terminal half-life (t_1/2_ in h) and the end time of the last sub-interval (t_n_ in h) were chosen based on the target PK profile to be simulated (**Table 2 and S2**)

**Table 2.**
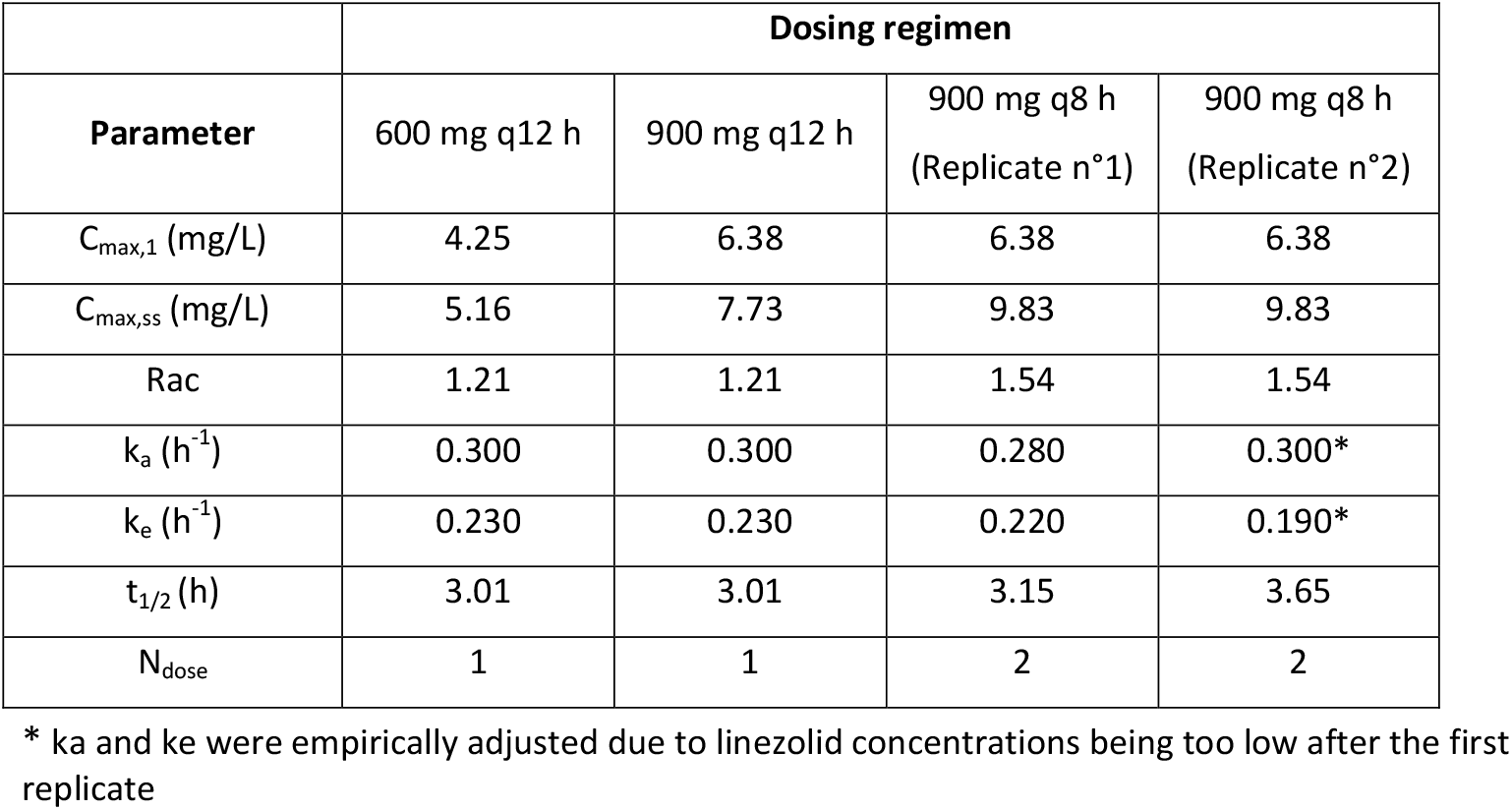
Target PK parameters for CSF concentrations.
2. The central reservoir + hollow-fiber cartridge volume (V_total_ in L) were chosen based on what was most practical in our experimental system. The number of sub-intervals (*n*) was chosen based on the capabilities of our programmable ambulatory infusion pump (**Table S2**).
3. The time to reach C_max,1_ was computed using equation (Eq.1).

| Parameter | Dosing regimen |  |  |  |
| --- | --- | --- | --- | --- |
|  | 600 mg q12 h | 900 mg q12 h | 900 mg q8 h<br>(Replicate n°1) | 900 mg q8 h<br>(Replicate n°2) |
| $C_{\text{max,1}}$ (mg/L) | 4.25 | 6.38 | 6.38 | 6.38 |
| $C_{\text{max,ss}}$ (mg/L) | 5.16 | 7.73 | 9.83 | 9.83 |
| Rac | 1.21 | 1.21 | 1.54 | 1.54 |
| $k_a$ ($\text{h}^{-1}$) | 0.300 | 0.300 | 0.280 | 0.300* |
| $k_e$ ($\text{h}^{-1}$) | 0.230 | 0.230 | 0.220 | 0.190* |
| $t_{1/2}$ (h) | 3.01 | 3.01 | 3.15 | 3.65 |
| $N_{\text{dose}}$ | 1 | 1 | 2 | 2 |

\* $k_a$ and $k_e$ were empirically adjusted due to linezolid concentrations being too low after the first replicate

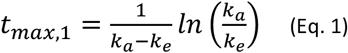
4. The dose administered at the end of the last sub-interval (Dose) with bioavailability (F) fixed to 1 and the fraction of target dose administered at the end of the last sub-interval (f_dose_) were computed using the equations (Eq. 2 and Eq. 3) below.

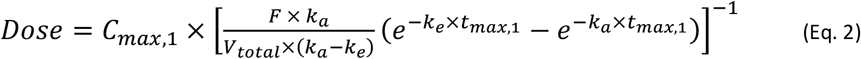

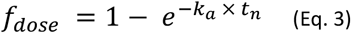

For each sub-interval i, t_i_ (h) the end time of sub-interval, A_i_ (mg) the amount of drug remaining to be administered for each t_i_ and S_i_ (mg/h) the infusion rate were computed using the equations (Eq. 4 to Eq. 6) below.

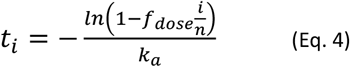

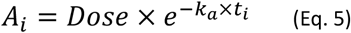

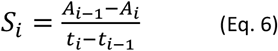

5. The linezolid concentration of the infused solution (C_infusion_) was chosen so that the volume administered during each sub-interval was as close as possible to 2 mL (lowest possible injectable volume with good precision with our ambulatory infusion pump) (**Table S2**).
6. For each sub-interval the infusion pump flow rate (Flow_infusion,i_) was computed using equation (Eq. 7) below:

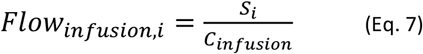
7. Infusion pump flow rates were then manually adjusted to the closest value programmable in the ambulatory infusion pump.

The flow of the pump adding diluent to the central reservoir (CL_elim_) was computed with equation (Eq. 8) (**Table S2**).

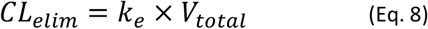

Diluent volume (V_diluent_) was then computed with equation (Eq. 9) for an experiment duration (Exp_duration_) of 96 h (**Table S2**).

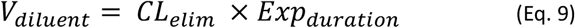

This algorithm is implemented in the freely accessible R-shiny application HF-App (25) and all experimental parameters used are summarized in the **Table S2**.

To ensure that linezolid unbound plasma and CSF concentrations were well reproduced in the HFIM, serial samples were collected from the central reservoir and the ECS of the cartridge at various time points. Samples were stored at −80°C until assayed by liquid chromatography coupled with tandem mass spectrometry (LC-MS/MS).

### 5. LC-MS/MS assay

Analysis of linezolid concentrations in CAMHB was performed by an LC-MS/MS method previously developed for assay of linezolid plasma concentrations (13). Calibration curves were established over 0.2 to 80 μg/mL. Samples were precipitated with 300 μL of acetonitrile containing the internal standard (Linezolid D8, purity of 100 % in powder form, Alsachim, Illkirch Graffenstaden, France) at 300 ng/mL and vortexed for 10 s. Then, samples were centrifuged at 14 000 rpm for 20 min at 4 °C and 100 μL of supernatant was transferred into a glass vial containing 100 μL of 10 mM ammonium formate. A volume of 2 μL was injected.

The system included a Shimadzu high-performance liquid chromatography system module (Nexera XR; Shimadzu, Marne la Vallée, France) coupled with a TQ3500 mass spectrometer (Sciex, Les Ulis, France). The compound was analyzed on an XBridge Peptide BEH300 C18 column (5 μm, 2.1 x 150 mm, Waters, Saint-Quentin-en-Yvelines, France). The mobile phase consisted of a mixture (50/50, v/v) of 10 mM ammonium formate and acetonitrile delivered isocratically at 0.20 mL/min. Electrospray ionization in positive mode was used for detection. Ions were analyzed in the multiple reaction monitoring, and the following transitions were inspected: m/z 338→296 and 346→304 for linezolid and internal standard, respectively.

The intraday variability was characterized at three concentrations levels (0.6, 15 and 60 μg/mL) with a precision < 3 % and bias < 9 %. The interday variability was characterized at three concentrations levels (0.6, 15 and 60 μg/mL) with a precision < 5 % and bias < 2 % (n = 19).

### 6. Modeling of observed concentrations in the HFIM reproducing plasma and CSF linezolid PK

#### Model structure

Linezolid concentrations observed in the central reservoir and ECS of the cartridge of the HFIM were analyzed simultaneously. A tri-compartmental model (**Figure 3**) was used to fit the data to account for a possible delay in linezolid concentrations in the ECS due to the diffusion between the ICS and ECS of the cartridge. Although it is assumed that the concentrations between the central reservoir and the cartridge equilibrate rapidly due to the high flow rate of the pump between the central reservoir and the cartridge, this has not been demonstrated (26,27).

**Figure 3.**
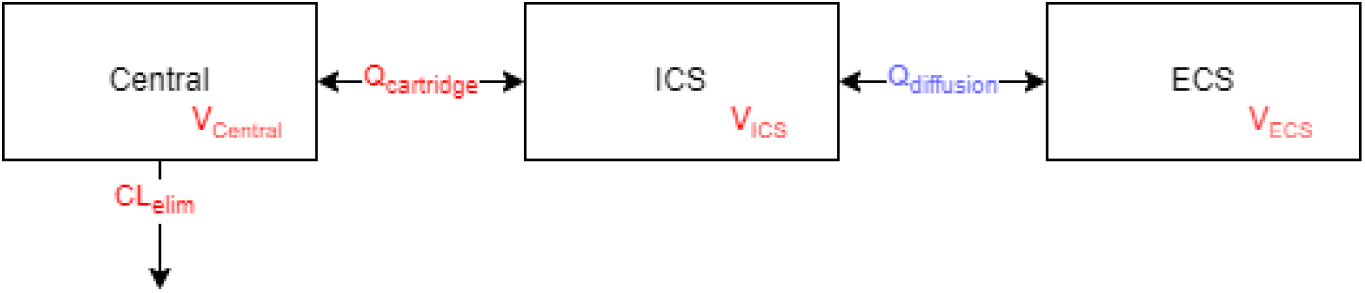
Structure of tri-compartmental model of HFIM. Central corresponds to the central reservoir of the HFIM, ICS to the intracapillary space of the cartridge, ECS to the extracapillary space of the cartridge, CL_elim_ to the pump flow rate from the central reservoir to the waste, Q_cartridge_ to the pump flow rate from the central reservoir to the cartridge and Q_diffusion_ to the diffusion flow rate between ICS and ECS of the cartridge. Red parameters were fixed, and blue parameter was estimated.

#### Estimation of constant diffusion flow rate between ICS and ECS

The tri-compartmental model allowed to estimate the constant diffusion flow rate between ICS and ECS of the cartridge (Q_diffusion_ in L/h) using Monolix (Monolix 2024R1, Lixoft SAS, a Simulations Plus company). To do so, the values of the central reservoir volume (V_Central_ in L), pump flow rate from the central reservoir to the waste (CL_elim_ in L/h), pump flow rate from the central reservoir to the cartridge (Q_cartridge_ in L/h) and the cartridge volume (V_cartridge_ in L) were fixed to experimental values (**Tables S1 and S2**). Volume of ICS (V_ICS_ in L) was computed by equation (Eq. 9) using the fraction of cartridge volume taken by fibers (F_ICS_, determined from the cartridge technical information given in **Text S2**) and the cartridge volume (V_cartridge_ in L). Volume of ECS (V_ECS,_ in L) was computed using equation (Eq. 10).

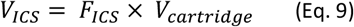

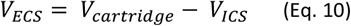

#### Validation of the tri-compartmental model

Model fit to data was evaluated by performing 1000 simulations with residual unexplained variability from the final model and plotting observed data against 90 % prediction interval. The model was considered valid when 90 % of observed data was within the 90 % prediction interval.

#### Validation of the HFIM setup

For each dosing regimen, the observed PK parameters in the ECS of the cartridge of the HFIM were computed by simulation under the final tri-compartmental model using the mrgsolve R package (23) with R software, version 4.2.2 (24). Computed observed PK parameters include:

- Maximal concentration after the first dose (C_max,1_) and at steady-state (C_max,ss_)
- Time to reach C_max_ after the first dose (T_max,1_) and at steady-state (T_max,ss_),
- Area under the curve for a dosing interval after the first dose (AUC_τ, 1_) and at steady state (AUC_τ, ss_)
- Elimination half-life (t_1/2_)

Our setup to reproduce plasma or CSF concentrations was considered valid when the computed observed PK parameters in the ECS of the cartridge were within 20 % of the target PK parameters.

## Results

### 1. Tri-compartmental model fits well the observed concentrations in the HFIM reproducing plasma and CSF linezolid PK

The tri-compartmental model fitted linezolid concentrations well, for the setup mimicking plasma (**Figure 4**) and CSF PK (**Figure 5**) for both the central reservoir and the ECS of the hollow-fiber cartridge.

**Figure 4.**
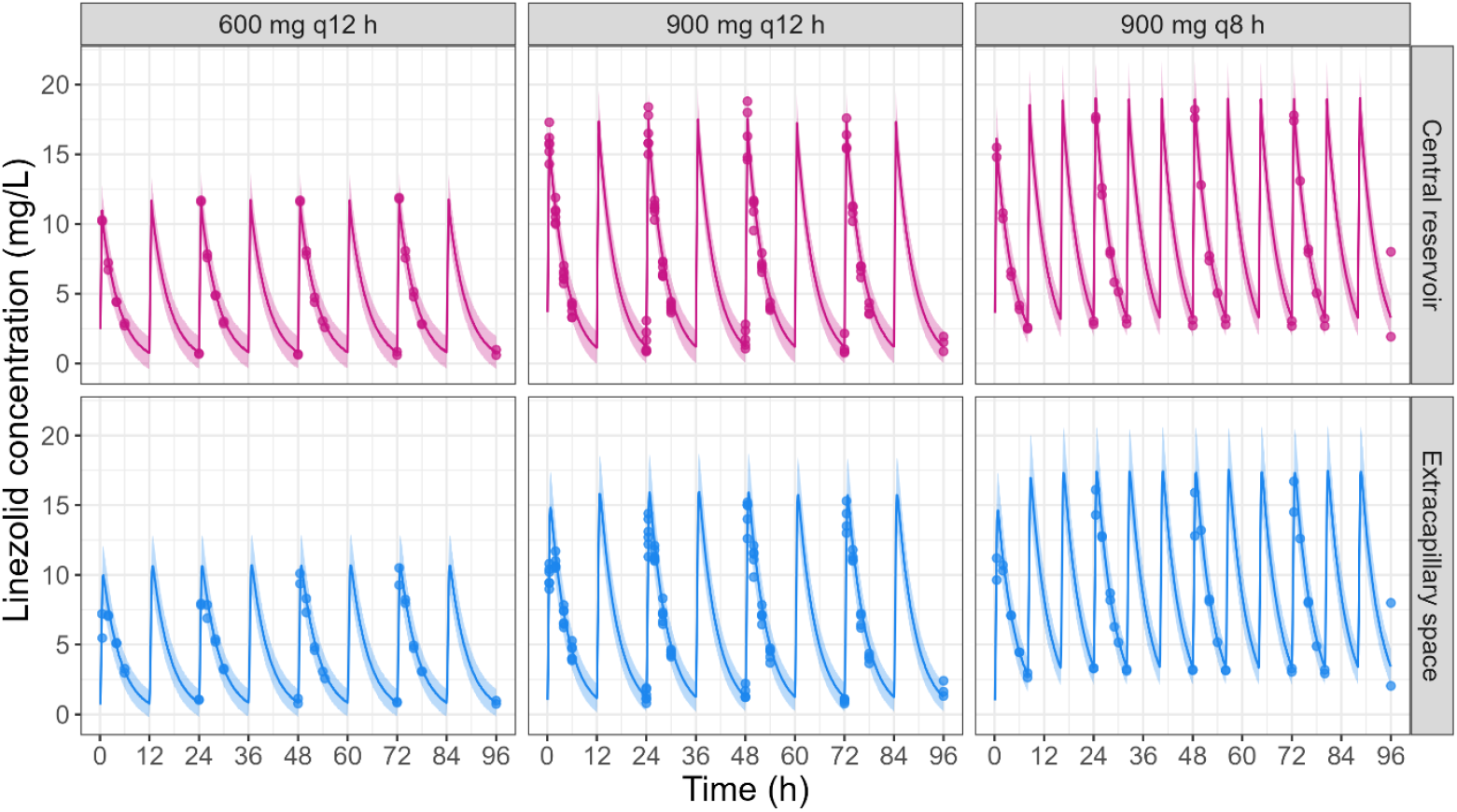
Observations versus simulations under the final model of linezolid concentrations observed in the central reservoir and in the cartridge ECS of the HFIM reproducing plasma PK for the three tested dosing regimens (*n* = 2 - 3). Points correspond to the observed linezolid concentrations in HFIM. The line corresponds to the median of the simulations. The colored area corresponds to the 90 % prediction interval (*i*.*e*. constructed from simulations with residual unexplained variability).

**Figure 5.**
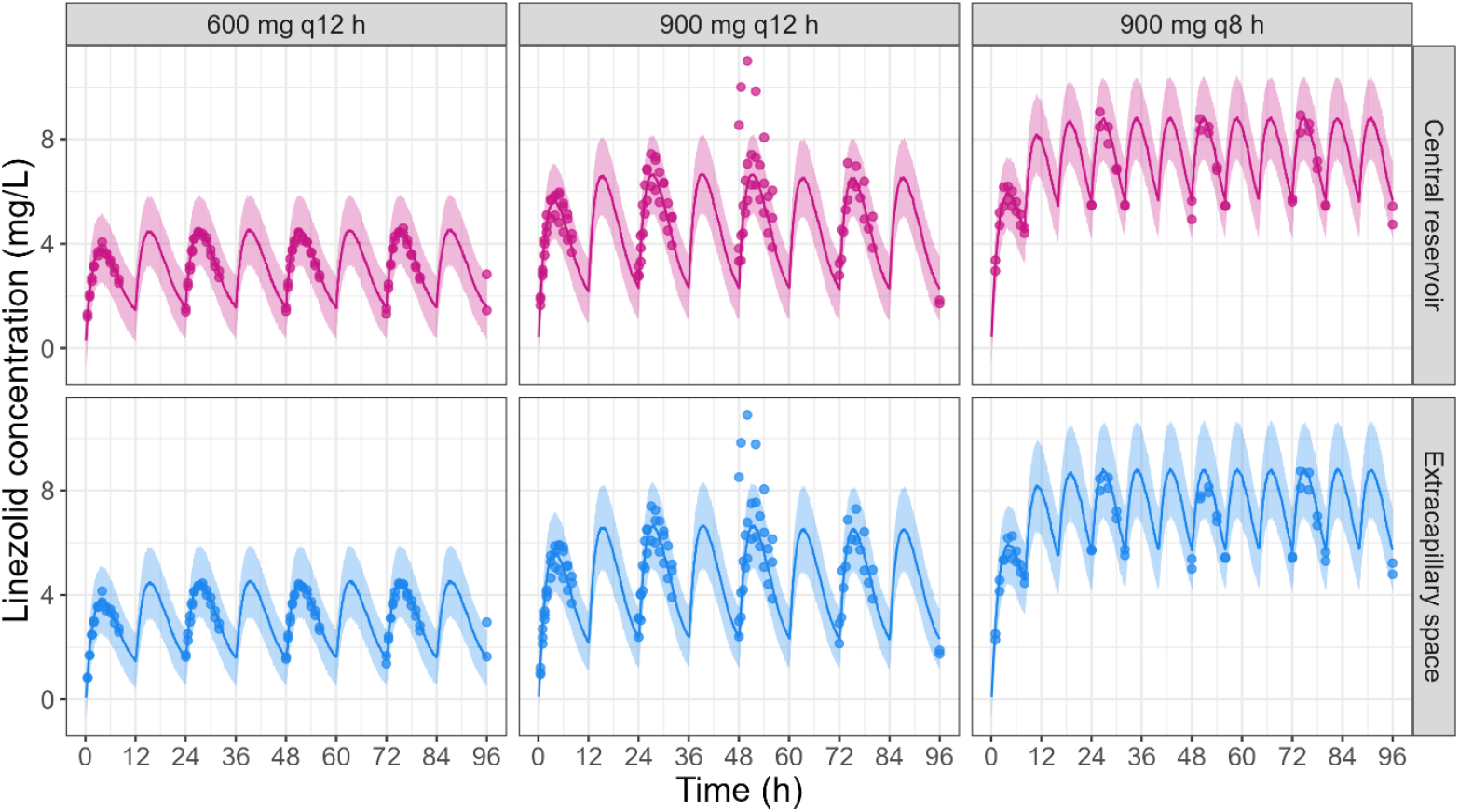
Observations versus simulations under the final model of linezolid concentrations observed in the central reservoir and in the cartridge ECS of the HFIM reproducing CSF PK for the three tested dosing regimens (*n* = 2 - 3). Points correspond to the observed linezolid concentrations in HFIM. The line corresponds to the median of the simulations. The colored area corresponds to the 90 % prediction interval (*i*.*e*. constructed from simulations with residual unexplained variability).

### 2. Diffusion across hollow-fiber membrane is quick but not instantaneous

The tri-compartmental model estimated a diffusion flow rate (Q_diffusion_) at 6.53 mL/min (0.392 L/h). Diffusion from the ICS to the ECS was not instantaneous, resulting in T_max_ in the cartridge ECS being delayed by 0.250 h when compared with the central reservoir concentrations when reproducing linezolid plasma concentrations whereas no difference was found when reproducing CSF concentrations (**Figure 6B**).

**Figure 6.**
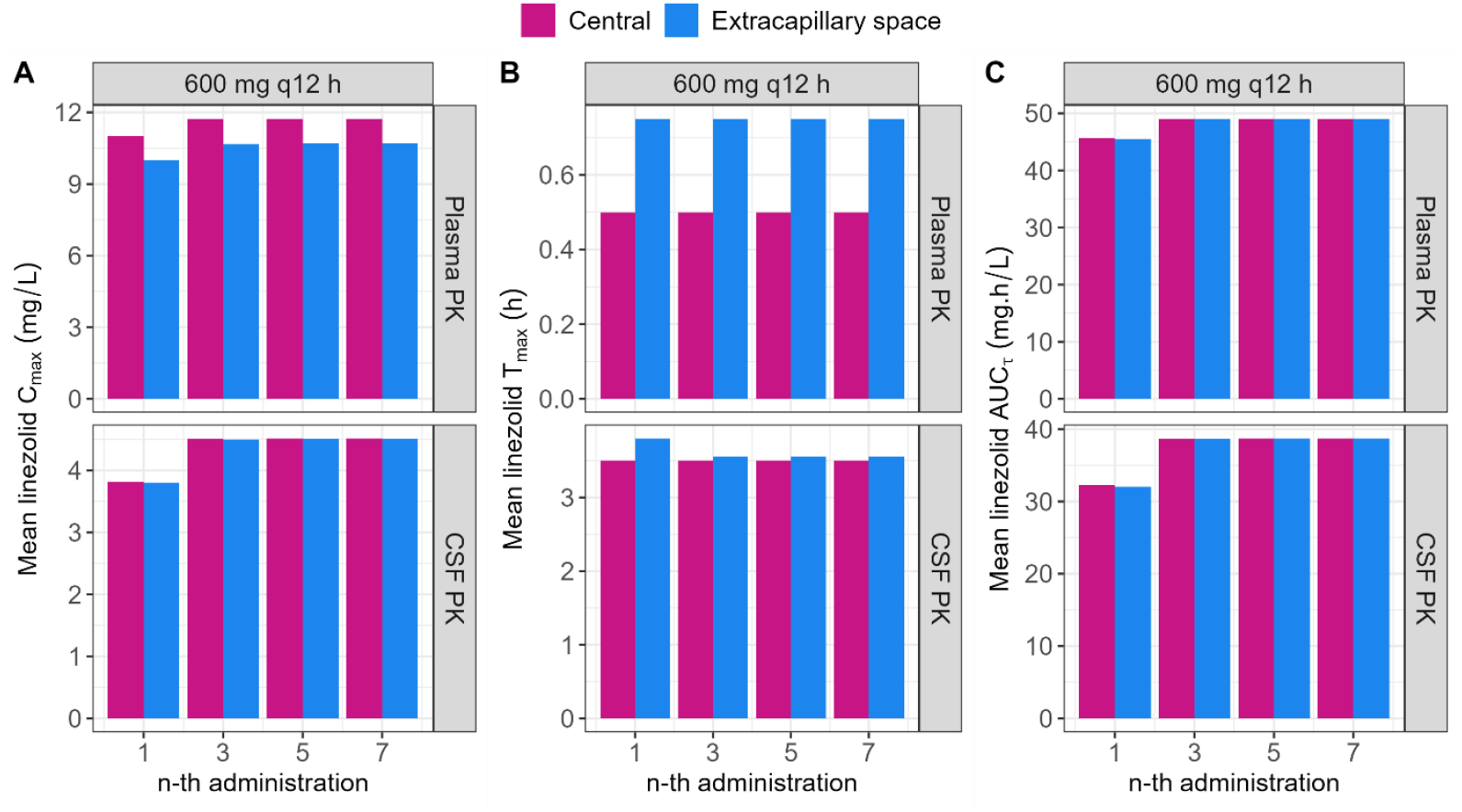
Linezolid PK parameters computed from observations in the central reservoir and in the cartridge ECS from HFIM experiments reproducing plasma PK (top panels) and CSF PK (bottom panels) after administration of 600 mg q12 h (*n* = 2). (A) Mean observed maximal concentrations (C_*max*_) of linezolid in the central reservoir (pink) and cartridge ECS (blue) after the 1^*st*^, 3^*rd*^, 5^*th*^ and 7^*th*^ administration. (B) Mean observed time to reach C_*max*_ (T_*max*_) of linezolid in the central reservoir (pink) and cartridge ECS (blue) after the 1^st^, 3^rd^, 5^th^ and 7^th^ administration. (C) Mean observed area under the curve over the dosing interval (AUC_τ_) of linezolid in the central reservoir (pink) and cartridge ECS (blue) after the 1^st^, 3^rd^, 5^th^ and 7^th^ administration.

When reproducing plasma concentrations, this delay prevented complete diffusion of linezolid to the cartridge ECS resulting in a ∼10 % lower observed C_max_ in the cartridge ECS when compared with the central reservoir concentrations, whereas no difference was found when reproducing CSF concentrations. (**Figure 6A**).

However, there was no impact on antibiotic exposure since the AUC_τ_ was the same between central reservoir and cartridge ECS for all simulated concentrations (**Figure 6C**).

Comparable results were observed for 900 mg q12 h and 900 mg q8 h dosing regimens (**Figure S1 and S2**).

### 3. Our Hollow-Fiber setup can be used to reproduce linezolid plasma PK

Target linezolid plasma concentrations from linezolid clinical population PK model (13) and observed linezolid concentrations in the cartridge ECS after an administration of 600 mg q12 h, 900 mg q12 h and 900 mg q8 h are shown in **Figure 7**.

**Figure 7.**
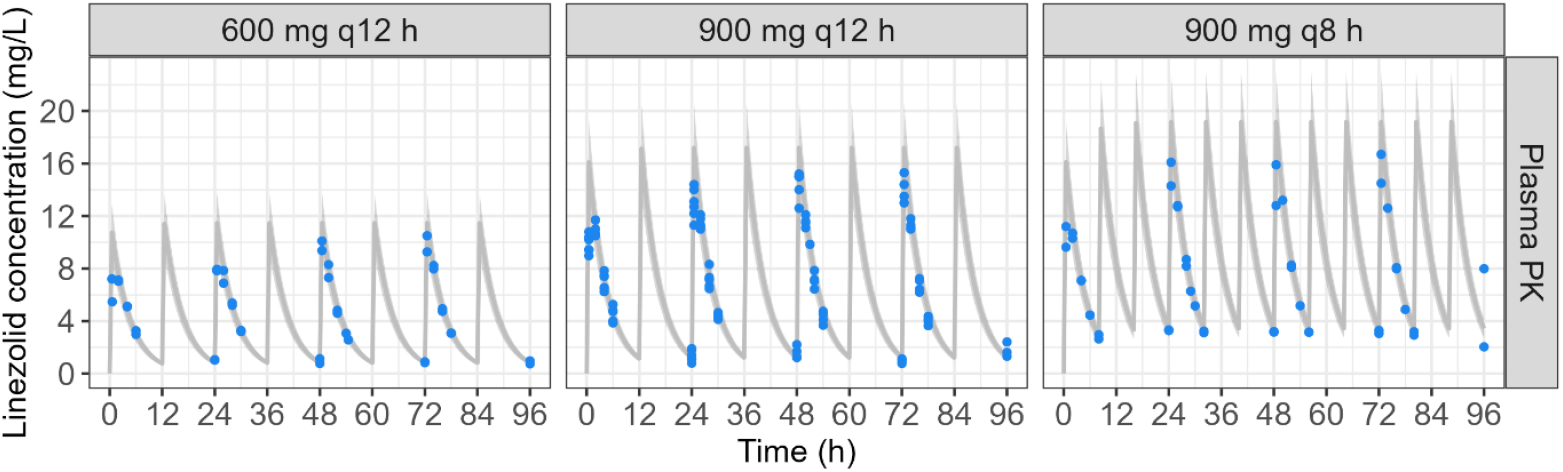
Target linezolid plasma concentrations from linezolid clinical population PK model (13) and observed linezolid concentrations in the cartridge ECS after infusion of 600 mg q12 h, 900 mg q12 h and 900 mg q8 h (*n* = 2 - 6). The gray line corresponds to target linezolid concentrations. The gray area corresponds to a 20 % bias from target linezolid concentrations. Points correspond to linezolid concentrations observed in the cartridge ECS of the HFIM.

Target and observed linezolid plasma PK parameters are compared in **Table 3**. C_max,1_, C_max,ss_, t_1/2_, AUC_τ, 1_ and AUC_τ, ss_ showed a bias lower than 10 %. T_max,1_ and T_max,ss_ absolute bias was 0.25 h (0.5 h versus 0.75 h) which we deemed acceptable.

**Table 3.** Target linezolid plasma PK parameters versus mean observed cartridge ECS linezolid PK parameters. *For T_max_, absolute bias (in h) was reported.

|  | Plasma |  |  |  |  |  |  |  |  |
| --- | --- | --- | --- | --- | --- | --- | --- | --- | --- |
|  | 600 mg q12 h |  |  | 900 mg q12 h |  |  | 900 mg q8 h |  |  |
|  | Target | ECS | Bias (%) | Target | ECS | Bias (%) | Target | ECS | Bias (%) |
| $C_{max,1}$ (mg/L) | 10.7 | 9.98 | -7.00 | 16.1 | 14.9 | -7.74 | 16.1 | 14.6 | -9.52 |
| $T_{max,1}$ (h) | 0.500 | 0.750 | 0.250* | 0.500 | 0.750 | 0.250* | 0.500 | 0.750 | 0.250* |
| $AUC_{\tau,1}$ (mgxh/L) | 46.1 | 45.6 | -1.23 | 69.2 | 67.8 | -2.02 | 61.6 | 58.9 | -4.28 |
| $C_{max,ss}$ (mg/L) | 11.5 | 10.7 | -6.81 | 17.2 | 15.9 | -7.55 | 19.2 | 17.4 | -9.04 |
| $T_{max,ss}$ (h) | 24.5 | 24.8 | 0.250* | 24.5 | 24.8 | 0.250* | 24.5 | 24.8 | 0.250* |
| $AUC_{\tau,ss}$ (mgxh/L) | 49.5 | 49.0 | -1.06 | 74.3 | 72.9 | -1.84 | 74.3 | 71.5 | -3.74 |
| $t_{1/2}$ (h) | 3.01 | 3.03 | 0.505 | 3.01 | 3.03 | 0.505 | 3.01 | 3.03 | 0.505 |

### 4. Our Hollow-Fiber setup can be used to reproduce linezolid CSF PK

The ambulatory infusion pump program to reproduce CSF concentrations after administration of 600 mg q12 h of linezolid is presented in **Table 4**. The corresponding expected PK profile after the first dose in the cartridge ECS is presented in **Figure 8** Error! Reference source not found.. Overall, the computed infusion pump program provided a good approximation of the target CSF concentrations. Comparable results were observed for the 900 mg q12 h and 900 mg q8 h dosing regimens (**Table S3 and Figure S3**).

**Table 4.**
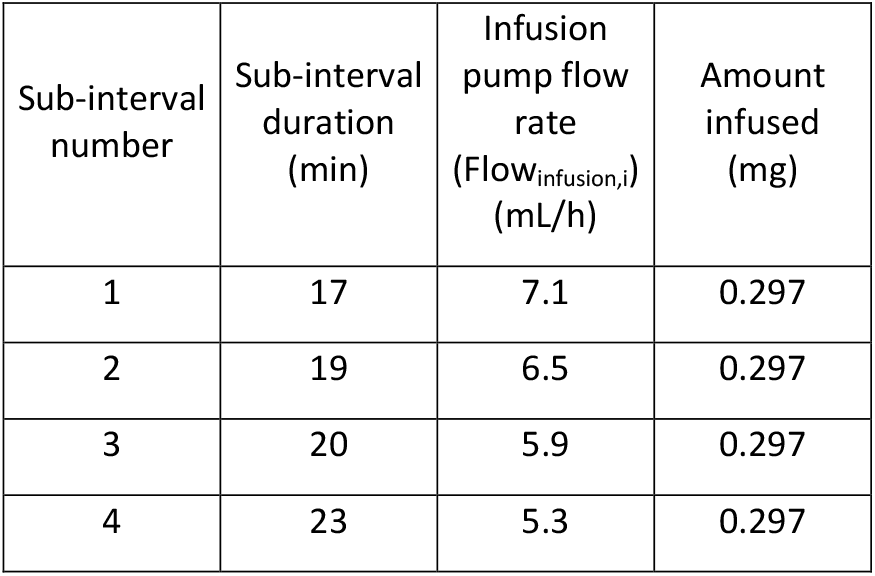
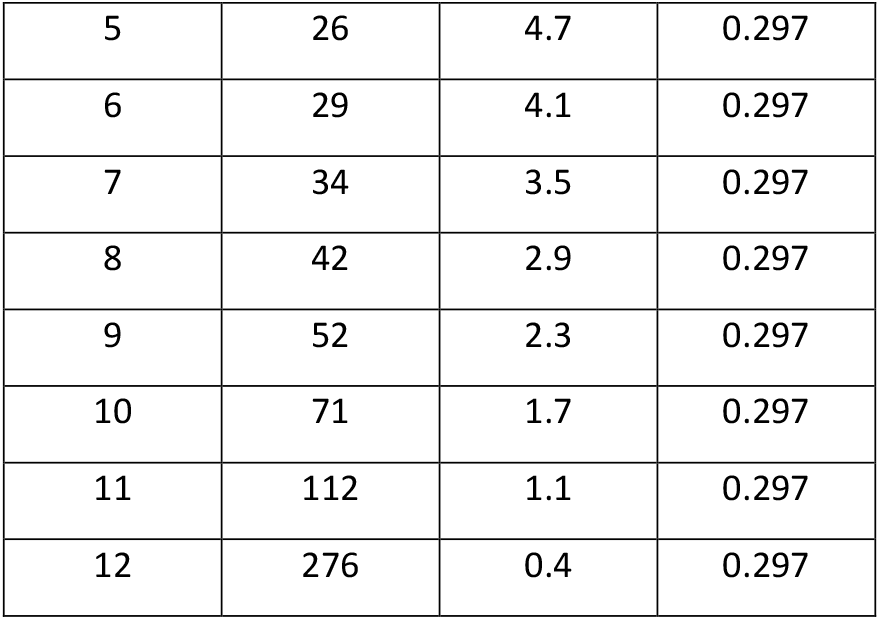
Ambulatory infusion pump program to reproduce CSF concentrations after administration of 600 mg q12 h of linezolid.

**Figure 8.**
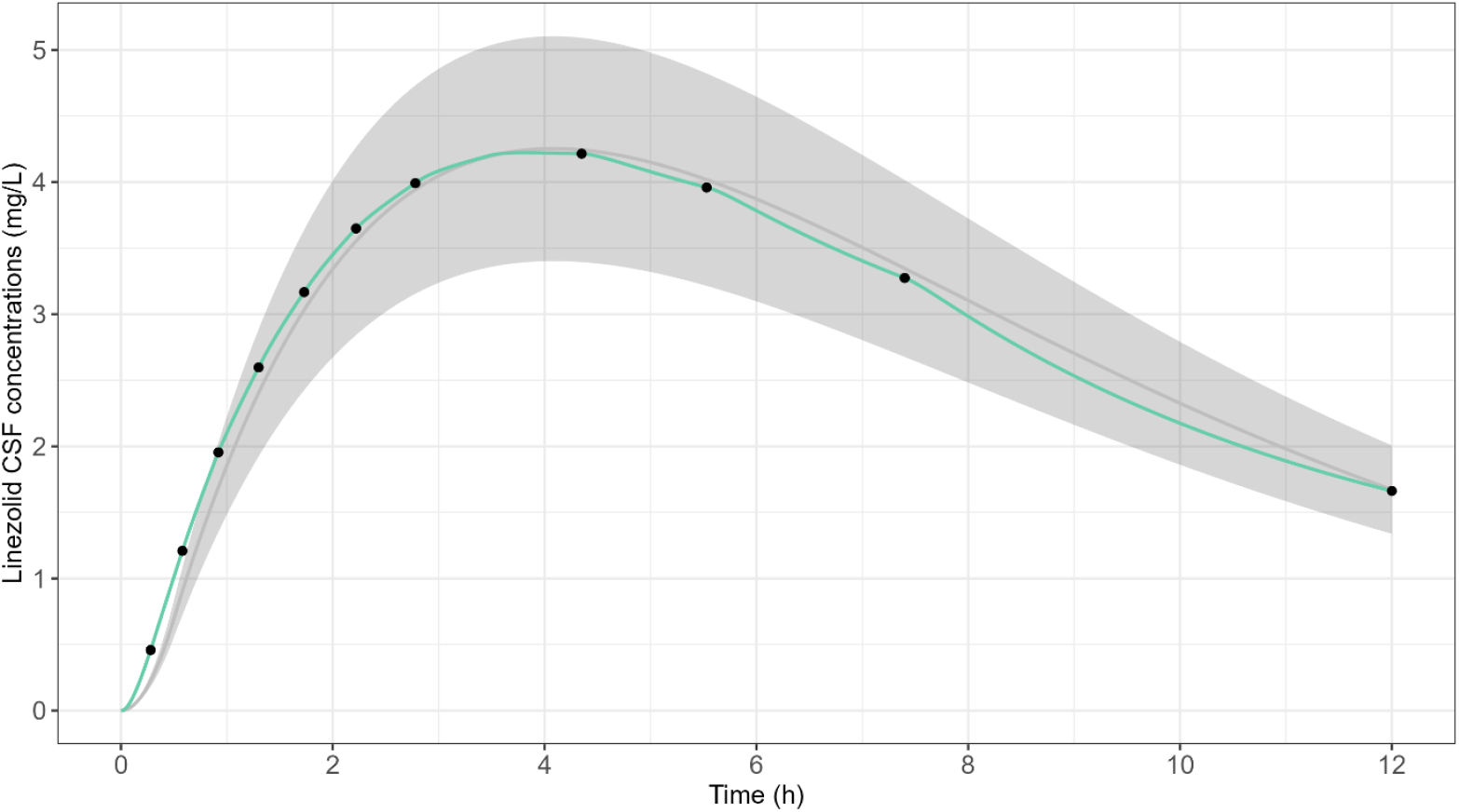
Target linezolid CSF concentrations from linezolid clinical population PK model (13) versus expected linezolid concentrations in the cartridge ECS for the first dose of 600 mg q12 h dosing regimen. The grey line corresponds to target linezolid concentrations. The grey area corresponds to a bias of 20 % from the target linezolid concentrations. The green line corresponds to expected linezolid concentrations in the cartridge ECS with the infusion pump program shown in **Table 4**. Black points correspond to expected concentrations at the end of each infusion sub-interval.

Target linezolid CSF concentrations from linezolid clinical population PK model (13) and observed linezolid concentrations in the cartridge ECS after an administration of 600 mg q12 h, 900 mg q12 h and 900 mg q8 h are shown in **Figure 9**.

**Figure 9.**
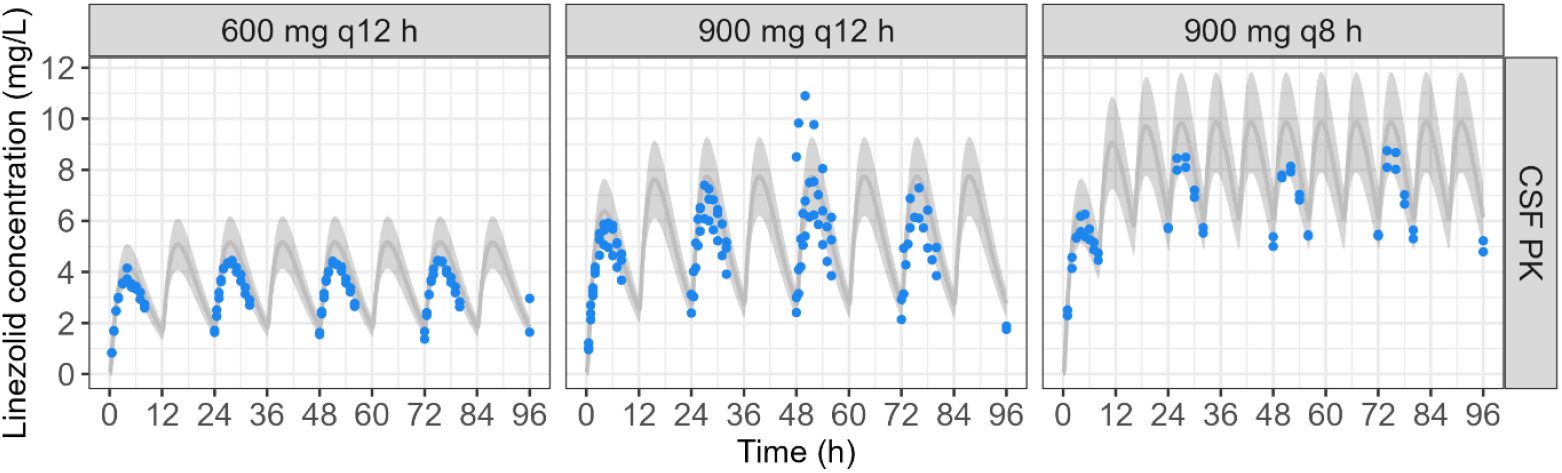
Target linezolid CSF concentrations from linezolid clinical population PK model (13) and observed linezolid concentrations in the cartridge ECS after infusion of 600 mg q12 h, 900 mg q12 h and 900 mg q8 h (*n* = 2 – 3). The gray line corresponds to target linezolid concentrations. The gray area corresponds to a 20 % bias from target linezolid concentrations. Points correspond to linezolid concentrations observed in the cartridge ECS of the HFIM.

Target and observed linezolid CSF PK parameters are compared in **Table 5**. C_max,1_, C_max,ss_, t_1/2_, AUC_τ, 1_ and AUC_τ, ss_ showed a bias lower than 20 %. T_max,1_ absolute bias was 0.3 h or less which we deemed acceptable. T_max,ss_ were unbiased.

**Table 5.** Target linezolid CSF PK parameters versus observed cartridge ECS linezolid PK parameters. *For T_max_, absolute bias (in h) was reported.

|  | 600 mg q12 h |  |  | 900 mg q12 h |  |  | 900 mg q8 h |  |  |  |  |  |
| --- | --- | --- | --- | --- | --- | --- | --- | --- | --- | --- | --- | --- |
|  |  |  |  |  |  |  | Replicate n°1 |  |  | Replicate n°2 |  |  |
|  | Target | ECS | Bias (%) | Target | ECS | Bias (%) | Target | ECS | Bias (%) | Target | ECS | Bias (%) |
| $C_{max,1}$ (mg/L) | 4.25 | 3.80 | -10.6 | 6.38 | 5.61 | -12.1 | 6.38 | 5.92 | -7.13 | 6.38 | 5.95 | -6.76 |
| $T_{max,1}$ (h) | 4.10 | 3.80 | -0.300* | 4.10 | 3.80 | -0.300* | 4.10 | 4.05 | -0.0500* | 4.10 | 4.00 | -0.100* |
| $AUC_{\tau,1}$ (mgxh/L) | 35.9 | 32.0 | -10.9 | 53.9 | 47.2 | -12.3 | 39.7 | 37.3 | -5.90 | 39.7 | 37.7 | -5.04 |
| $C_{max,ss}$ (mg/L) | 5.16 | 4.50 | -12.6 | 7.73 | 6.64 | -14.1 | 9.83 | 8.50 | -13.5 | 9.83 | 9.04 | -8.07 |
| $T_{max,ss}$ (h) | 27.6 | 27.6 | 0.000* | 27.6 | 27.6 | 0.000* | 27.0 | 27.0 | 0.000* | 27.0 | 26.9 | -0.0500* |
| $AUC_{\tau,ss}$ (mgxh/L) | 44.7 | 38.7 | -13.4 | 67.0 | 57.1 | -14.8 | 66.9 | 58.4 | -12.7 | 66.9 | 62.8 | -6.13 |
| $t_{1/2}$ (h) | 3.01 | 3.03 | 0.505 | 3.01 | 3.03 | 0.505 | 3.15 | 3.16 | 0.387 | 3.65 | 3.66 | 0.264 |

## Discussion

Most HFIM studies found in the literature assay drug concentrations in the central reservoir rather than in the ECS (4,5,7–9,28–32). However, since bacteria are trapped within the ECS, it represents the true site of action, thus concentrations in the ECS are the most relevant to PK/PD studies. The reason why most studies do not assay drugs in the ECS is that it is usually assumed that concentrations in the ECS are in very rapid equilibrium with the central reservoir concentrations.

Some studies, challenged this assumption by measuring concentrations in both the central reservoir and ECS and reported that a delay of 15 min (6) and 20 min (18) was necessary to reach equilibrium between central reservoir and the ECS of the cartridge after short infusions of 30 min or 1 h.

In line with those previous reports, we also observed a delay of 15 min (0.25 h) in reaching equilibrium between the central reservoir and ECS when we reproduced 30 min short infusions of linezolid (**Figure 6B**). However, when reproducing CSF PK profiles, no such delay was observed (**Figure 6B**). Our hypothesis is that this delay is proportional to the rate of change in central reservoir concentrations, the higher the infusion rate and the lower the half-life, the longer the time to equilibrium will be.

This delay not only has an impact on the time necessary to reach maximal concentration (T_max_) in the ECS but also on the maximal concentration itself (C_max_). Indeed, while the drug concentrations equilibrate between the central reservoir and the ECS, the drug is also eliminated from the central reservoir, reducing the overall amount of drug that reaches the ECS. In our study, the impact on C_max_ was limited (< 10 % **Table 3**). However, if the delay is truly proportional to infusion rate and half-life, there will be future cases where the impact of C_max_ will be significant.

In such cases, the parameterization of the HFIM could be adjusted in order to take into account this equilibration delay. In order to do so, one would have to first estimate the rate of diffusion from the ICS to the ECS (Q_diffusion_) and then perform numerical optimization to find the best HFIM parameters to reproduce the desired concentration time-curve.

Our results further reinforce the need to measure ECS concentrations when performing HFIM experiments when one wants to study the PK/PD of an anti-infective drug. Furthermore, it is especially important for antibiotics whose physicochemical properties render susceptible to sticking to hollow-fibers (26) or to bacteria-mediated degradation (e.g., β-lactamase-producing strains) (33).

A limitation of our validation is that when we computed the target PK parameters to reproduce linezolid CSF concentrations in the HFIM, we empirically adjusted the fitted ke and ka values to yield a better match between the expected concentrations and the target concentrations. We made a mistake when performing this step for dosing regimens 600 mg q12 h, 900 mg q12 h and the first replicate of 900 mg q8 h where we set ke to 0.23 h^-1^ or 0.22 h^-1^ instead of 0.19 h^-1^ (**Table S2**). This resulted in faster linezolid elimination and therefore in lower C_max_ and AUC_τ_.

However, in spite of the mistake, bias of the observed PK parameters remained lower than 20 % for all tested dosing regimens (*n* = 3), target PK (*n* = 2 per dosing regimen), and all replicates (*n* ≥ 2 for each setup) (**Table 3 and Table 5**).

We developed and validated a new HFIM setup that enables *in vitro* reproduction of mono-compartmental PK with an absorption phase. This setup was applied to successfully reproduce site of action PK of linezolid, but could also be used to reproduce plasma PK after oral administration. An important strength of our setup is that it does not come with an overly complex modification to the material setup of the experiment, adding only a programmable infusion pump to the system (as opposed to adding a whole compartment and peristaltic pump).

This was made possible by the development of an algorithm that computes infusion rates and durations that enable an approximation of the first-order absorption kinetics by a series of continuous infusions. Since manual application of the algorithm would be tedious, we included it in an open-source web application designed to help design experimental protocols for HFIM (25). Thus, our setup and algorithm are easy to translate to any other HFIM experiment.

## Acknowledgment

The Shimadzu high-performance liquid chromatography system module (Nexera XR; Shimadzu, Marne la Vallée, France) coupled with a TQ3500 mass spectrometer (Sciex, Les Ulis, France) was obtained through the support of the Nouvelle Aquitaine CPER and FEDER programs.

## Data availability statement

The data used in this study are available at the following address https://doi.org/10.57745/NRXPOP.

## Supplementary data

**Table S1:** Experimental parameters used in HFIM to simulate plasma concentrations.

|  | Parameter | Parameter description | 600 mg q12 h | 900 mg q12 h | 900 mg q8 h |
| --- | --- | --- | --- | --- | --- |
| Set in the application | $V_{\text{Central}}$ (L) | Volume in the central reservoir | 0.300 | 0.300 | 0.300 |
| | $Q_{\text{Cartridge}}$ (L/h) | Pump flow rate from the central reservoir to the cartridge | 3.60 | 3.60 | 3.60 |
| | $V_{\text{Cartridge}}$ (L) | Volume of the cartridge | 0.0600 | 0.0600 | 0.0600 |
| | $t_{\text{infusion}}$ (h) | Infusion duration | 0.500 | 0.500 | 0.500 |
| | $V_{\text{infusion}}$ (mL) | Volume of infusion | 5.00 | 5.00 | 5.00 |
| | $\tau$ (h) | Dosing interval | 12 | 12 | 8 |
| | $n_{\text{doses}}$ | Total number of doses | 8 | 8 | 12 |
| | $\text{Exp}_{\text{duration}}$ (h) | Experiment duration | 96 | 96 | 96 |
| | $C_{\text{max},1}$ (mg/L) | Maximal concentration after the first dose | 10.7 | 16.1 | 16.1 |
| | $t_{1/2}$ (h) | Terminal half-life | 3.01 | 3.01 | 3.01 |
| Computed by the application | $CL_{\text{elim}}$ (L/h) | Pump flow rate from the central reservoir to the waste | 0.0828 | 0.0828 | 0.0828 |
| | $V_{\text{diluent}}$ (L) | Volume of diluent | 7.95 | 7.95 | 7.95 |
| | $C_{\text{infusion}}$ (mg/L) | Infusion solution concentration | 838 | 1220 | 1230 |
| Performed | Replicates | Number of replicates | 2 | 6 | 2 |

**Table S2:** Experimental parameters used in HFIM to simulate CSF concentrations.

|  | Parameter | Parameter description | 600 mg q12<br>h | 900 mg q12<br>h | 900 mg q8 h<br>(n°1) | 900 mg q8 h<br>(n°2) |
| --- | --- | --- | --- | --- | --- | --- |
| Set in the application | V<sub>Central</sub> (L) | Volume in the central reservoir | 0.300 | 0.300 | 0.300 | 0.300 |
|  | Q<sub>Cartridge</sub> (L/h) | Pump flow rate from the central reservoir to the cartridge | 3.60 | 3.60 | 3.60 | 3.60 |
|  | V<sub>Cartridge</sub> (L) | Volume of the cartridge | 0.0600 | 0.0600 | 0.0600 | 0.0600 |
|  | t<sub>1/2</sub> (h) | Terminal half-life | 3.01 | 3.01 | 3.15 | 3.65* |
|  | F | Bioavailability | 1 | 1 | 1 | 1 |
|  | k<sub>a</sub> (h<sup>-1</sup>) | Absorption rate constant | 0.300 | 0.300 | 0.280 | 0.300* |
|  | C<sub>max, 1</sub> (mg/L) | Maximal concentration after the first dose | 4.25 | 6.38 | 6.38 | 6.38 |
|  | Exp<sub>duration</sub> (h) | Experiment duration | 96 | 96 | 96 | 96 |
|  | n | Number of sub-intervals | 12 | 12 | 8 | 8 |
|  | t<sub>n</sub> (h) | The end time of the last sub-interval | 12 | 12 | 8 | 8 |
|  | n<sub>doses</sub> | Total number of doses | 8 | 8 | 12 | 12 |
|  | V<sub>infusion</sub> (mL) | Volume of infusion | 2.00 | 2.00 | 3.00 | 2.00 |
| Computed by the application | t<sub>max,1</sub> (h) | Time to reach C<sub>max,1</sub> | 3.80 | 3.80 | 4.02 | 4.15 |
|  | Dose (mg) | Dose administered at the end of the last sub-interval | 3.66 | 5.50 | 5.56 | 5.06 |
|  | f<sub>dose</sub> | Fraction of target dose administered at the end of the last sub-interval | 0.973 | 0.973 | 0.894 | 0.909 |
|  | CL<sub>elim</sub> (L/h) | Pump flow rate from the central reservoir to the waste | 0.0828 | 0.0828 | 0.0792 | 0.0684 |
|  | V<sub>diluent</sub> (L) | Volume of diluent | 7.95 | 7.95 | 7.60 | 6.57 |
|  | C<sub>infusion</sub> (mg/L) | Infusion solution concentration | 148 | 222 | 207 | 288 |
| Performed | Replicates | Number of replicates | 2 | 3 | 1 | 1 |

\* ka and ke were empirically adjusted due to linezolid concentrations being too low after the first replicate

### 3. Text S1: Detailed explanation of our mathematical algorithm

#### Objective

Find an experimental setup to be able to simulate first-order absorption and elimination within a hollow-fiber experiment.

#### Setup

Use the standard two-compartment hollow-fiber system (central reservoir + hollow fiber cartridge) with a variable flow infusion to simulate the drug absorption

#### Challenges

The challenge with the aforementioned setup is to find a procedure to compute the infusion parameters (*i*.*e*. concentration, volume and flow) with a focus on how often and how to change the infusion flow over time.

#### Theoretical solution

An analytical formula giving the infusion flow rate (*I*(*t*)) necessary to maintain a constant absorption rate parameter (*k*_*a*_(*t*)) exists. It can be derived as follows:

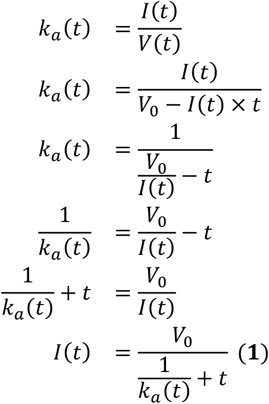

Where: *t* is the time post beginning of the infusion. *V*(*t*) is the volume of drug solution remaining in the infusion bag at time *t. V*_0_ is the initial volume in the infusion bag.

This equation is solvable for a known constant *k*_*a*_(*t*) and *V*_0_.

However, this solution is not applicable in our case because, we do not have access to a pump that allows for programming of a continuously varying infusion rate thus cannot apply this formula directly and instead need to approximate it by multiple discrete infusion rates changing at discrete times.

#### Practical solution

Let *n* be the number of discrete infusions rates that we will use in the experiment. Let *t*_*i*_ be the end time of sub-interval *i* where *t* ∈ [*t*_*i*−1_, *t*_*i*_ [ and *t*_0_ = 0 and *A*_*i*_ be the amount of drug remaining to be administered for each *t*_*i*_.

We need to define the duration of the dosing sub-intervals. We can define them by splitting the total dosing duration into *n* sub*-*intervals in such a way that the amount of drug infused to the central reservoir during each sub-interval is equal. Mathematically:

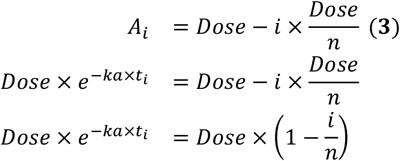

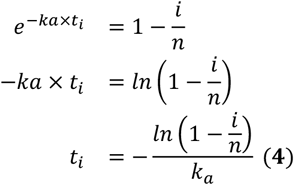

However, in this setup when *i* = *n, t*_*i*_ = +∞ which is longer than any sane experimentalist is willing to wait.

There are two ways to solve this issue. We ask the experimentalist to either define a fraction of the total dose that they want to be administered at the end of the last sub-interval, or specify the time at which the last sub-interval should end, in which case we will return the % of total dose administered by then. In this paper, since we are dealing with repeated doses with a predefined dosing interval, we chose the latter.

Let *f*_*dose*_ be the fraction of total dose administered at the end of the last sub-interval (*t*_*n*_). We can transform equation (**3**) to have *f*_*dose*_ and *t*_*n*_

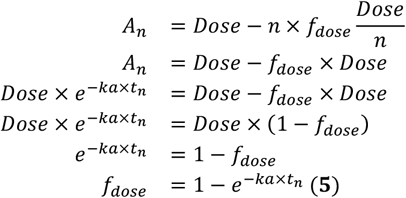

And in this case equation (**4**) has to be modified to take into account this forced end of last sub-interval while maintaining even sub-intervals with regards to amount transferred to central reservoir.

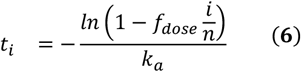

To put our solution into practice, we start by defining the time at which our last sub-interval will end (*t*_*n*_), then we can use it in equation (**5**) to compute *f*_*dose*_. Afterwards, we can use equation (**6**) to compute all *t*_*i*_.

Then we can use the standard pharmacokinetic equation for first-order absorption to compute *A*_*i*_ defined as:

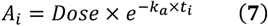

We can then simply compute the slope of the *A(t)* curve for each sub-interval and use this as our infusion rate, we will call it *S*_*i*_:

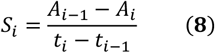

This will allow *A*_*i*_ to be equal to what it would have been with a first-order absorption process at the end of each sub-interval.

Bringing it all together we can write our algorithm in pseudocode as follows:

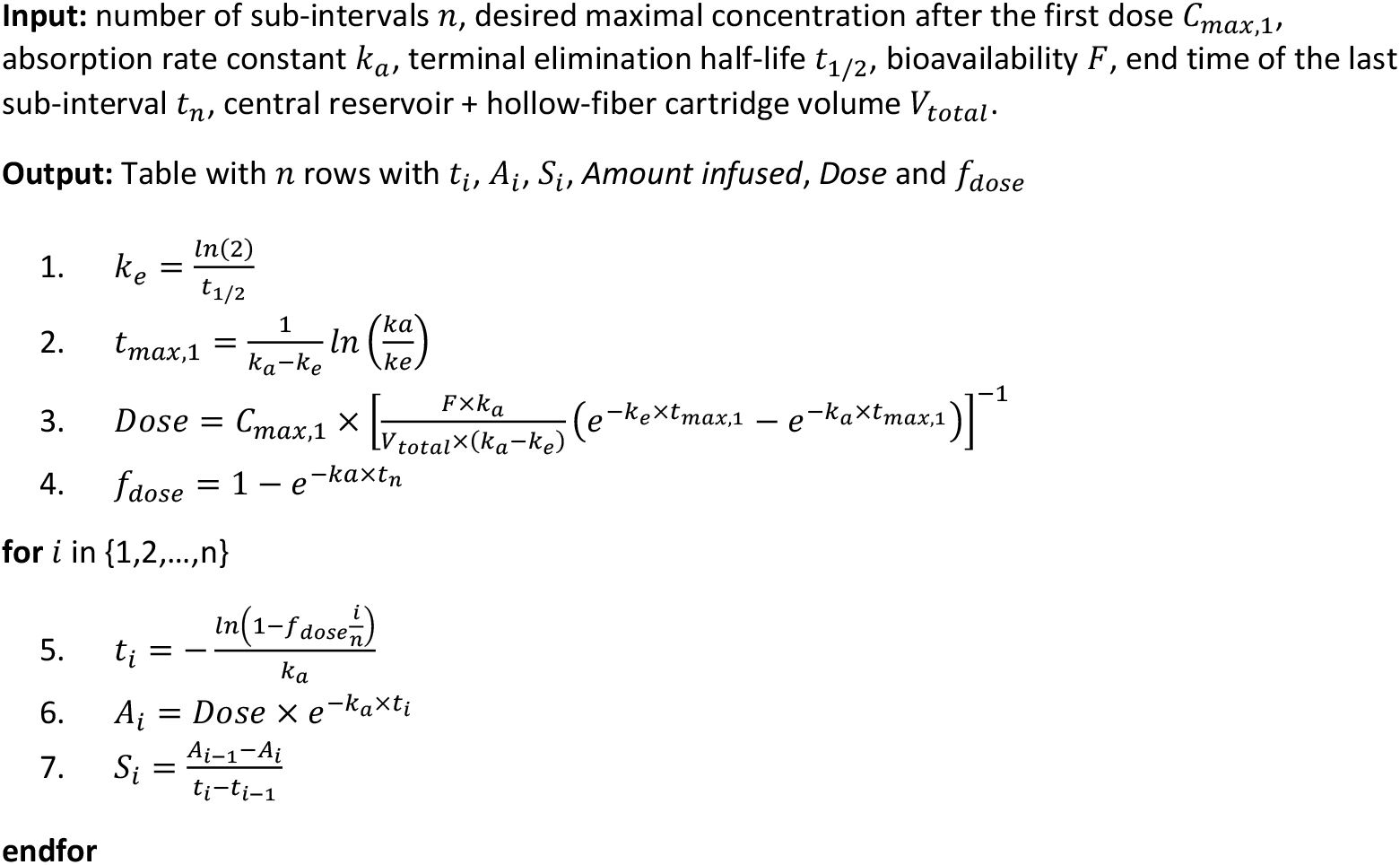

**Example case**

*n* = 12, *C*_*max*_ = 4.25*mg/L, k*_*a*_=0.36*h*^*−*^*1, t*_1/2_ = 3.15*h, V*_*total*_ = 0.360*L, C* = 1000*mg/L, F = 1, t*_*n*_ = 12*h*

| $t_i$ (h) | $A_i$ (mg) | $S_i$ (mg/h) | *Amount infused* (mg) |
| --- | --- | --- | --- |
| 0.00 | 3.32 | NA | NA |
| 0.24 | 3.04 | 1.14 | 0.27 |
| 0.50 | 2.77 | 1.05 | 0.27 |
| 0.79 | 2.50 | 0.95 | 0.27 |
| 1.11 | 2.23 | 0.85 | 0.27 |
| 1.47 | 1.95 | 0.75 | 0.27 |
| 1.89 | 1.68 | 0.65 | 0.27 |
| 2.38 | 1.41 | 0.55 | 0.27 |
| 2.98 | 1.14 | 0.46 | 0.27 |
| 3.74 | 0.86 | 0.36 | 0.27 |
| 4.80 | 0.59 | 0.26 | 0.27 |
| 6.52 | 0.32 | 0.16 | 0.27 |
| 12.00 | 0.04 | 0.05 | 0.27 |

*Dose = 3*.3 mg *f*_*dose*_ = 98.7 %

### 4. Text S2: Determination of fraction of fibers in the cartridge

Our goal is to compute the fraction of the total cartridge volume (F_ICS_) that is occupied by hollow-fibers (FX paed helixone dialyzer, Fresenius Medical Care, Bad Homburg, Germany).

From the cartridge documentation, we have:

| Fiber lumen diameter (cm) | Fiber wall thickness (cm) | Effective surface area (cm<sup>2</sup>) |
| --- | --- | --- |
| 0.0220 | 0.00350 | 2000 |

From direct measurements of the cartridge, we have:

| Cartridge length (cm) | Cartridge perimeter (cm) |
| --- | --- |
| 25.0 | 6.00 |

We will assume that the cartridge and the fibers are straight cylinders.

We can compute the volume of an individual fiber (V_fiber_):

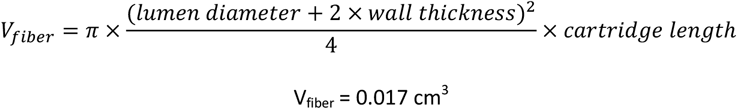

We can also compute the surface area of 1 fiber (SA_fiber_):

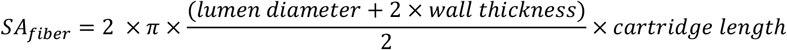

Note that we omit the surface area of the horizontal cross section of the cylinder since it is unavailable for drug diffusion.

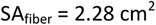

In the cartridge documentation, we are provided with the effective surface area of the hollow fibers which is the sum of the surface area of all the hollow fibers. We can thus deduce the number of fibers in the cartridge n_fibers_:

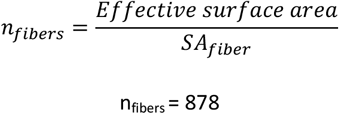

We can then compute the total volume occupied by the fibers (V_fibers,total_) by multiplying V_fiber_ by n_fibers_:

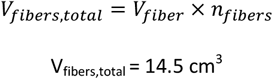

From the perimeter of the cartridge, we can compute the diameter of the cartridge (d_cartridge_):

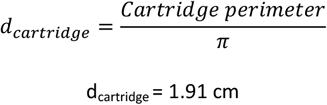

We can then compute the theoretical volume of the cartridge:

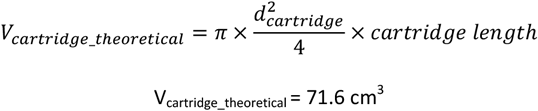

Finally, we can compute the fraction of cartridge volume occupied by the fibers:

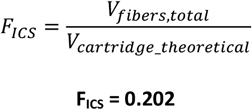

**Figure S1:**
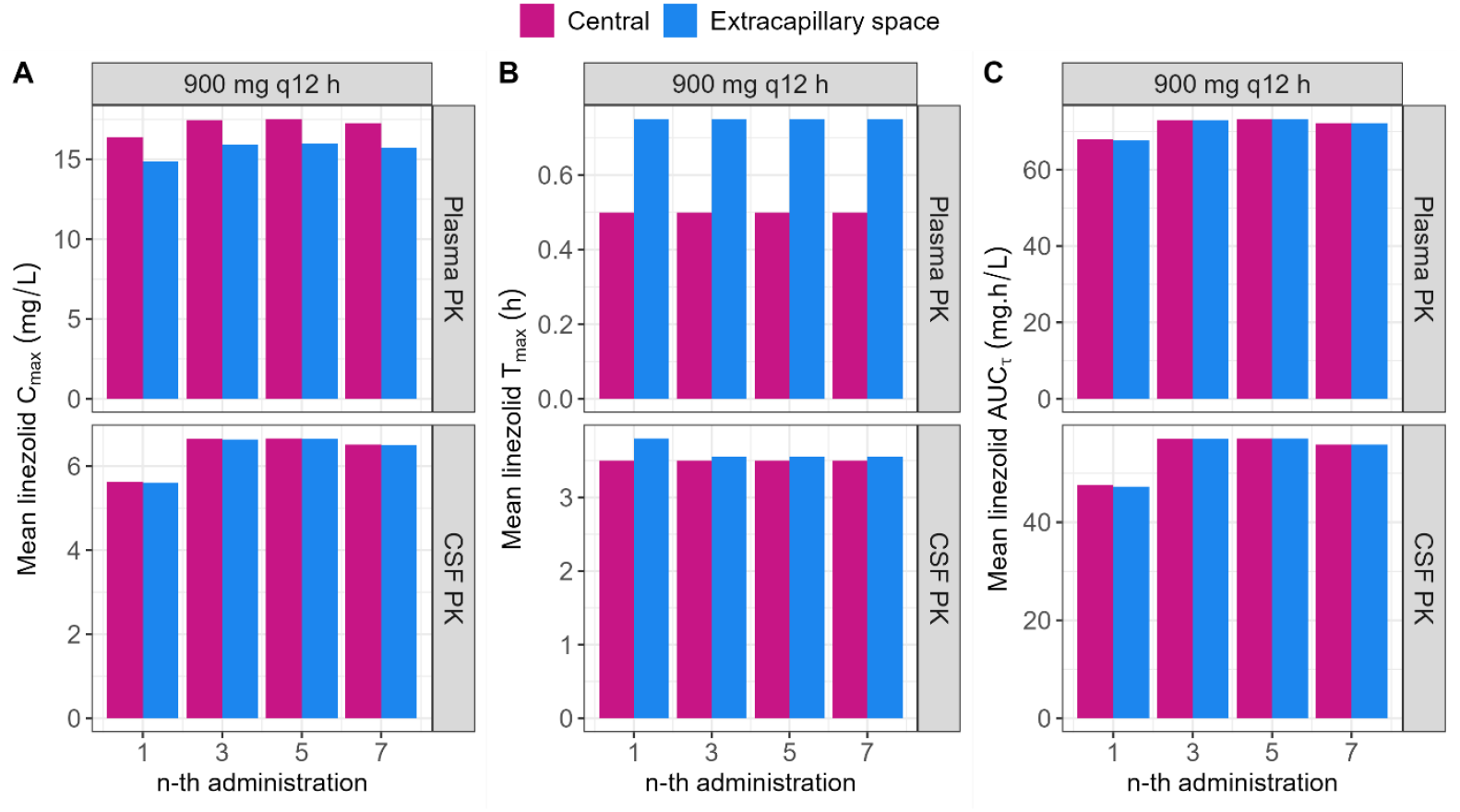
Linezolid PK parameters computed from observations in the central reservoir and in the cartridge ECS from HFIM experiments reproducing plasma PK (top panels) and CSF PK (bottom panels) after administration of 900 mg q12 h (*n* = 2-6). (A) Mean observed maximal concentrations (C_*max*_) of linezolid in the central reservoir (pink) and cartridge ECS (blue) after the 1^*st*^, 3^*rd*^, 5^*th*^ and 7^*th*^ administration. (B) Mean observed time to reach C_*max*_ (T_*max*_) of linezolid in the central reservoir (pink) and cartridge ECS (blue) after the 1^st^, 3^rd^, 5^th^ and 7^th^ administration. (C) Mean observed area under the curve over the dosing interval (AUC_τ_) of linezolid in the central reservoir (pink) and cartridge ECS (blue) after the 1^st^, 3^rd^, 5^th^ and 7^th^ administration.

**Figure S2:**
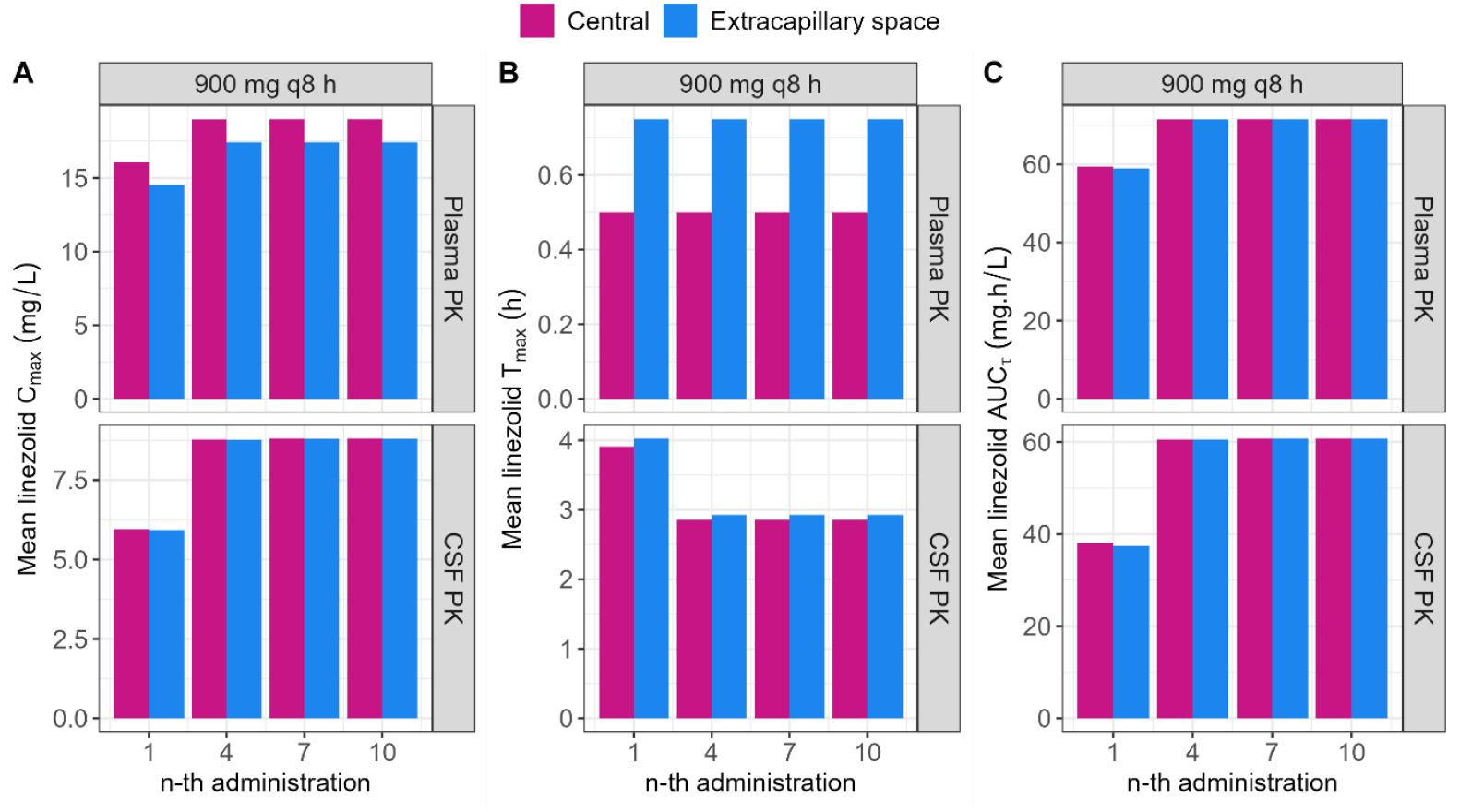
Linezolid PK parameters computed from observations in the central reservoir and in the cartridge ECS from HFIM experiments reproducing plasma PK (top panels) and CSF PK (bottom panels) after administration of 900 mg q8 h (*n* = 2). (A) Mean observed maximal concentrations (C_*max*_) of linezolid in the central reservoir (pink) and cartridge ECS (blue) after the 1^*st*^, 4^*th*^, 7^*th*^ and 10^*th*^ administration. (B) Mean observed time to reach C_*max*_ (T_*max*_) of linezolid in the central reservoir (pink) and cartridge ECS (blue) after the 1^*st*^, 4^*th*^, 7^*th*^ and 10^*th*^ administration. (C) Mean observed area under the curve over the dosing interval (AUC_τ_) of linezolid in the central reservoir (pink) and cartridge ECS (blue) after the 1^*st*^, 4^*th*^, 7^*th*^ and 10^*th*^ administration.

**Table S3:** Ambulatory infusion pump program to reproduce CSF concentrations after administration of 900 mg q12 h and 900 mg q 8 h of linezolid.

| 900 mg q12 h |  |  |  | 900 mg q 8h (Replicate n°1) |  |  |  | 900 mg q8 h (Replicate n°2)* |  |  |  |
| --- | --- | --- | --- | --- | --- | --- | --- | --- | --- | --- | --- |
| Sub-interval number | Sub-interval duration (min) | Infusion pump flow rate (Flow<sub>infusion,i</sub>) (mL/h) | Amount infused (mg) | Sub-interval number | Sub-interval duration (min) | Infusion pump flow rate (Flow<sub>infusion,i</sub>) (mL/h) | Amount infused (mg) | Sub-interval number | Sub-interval duration (min) | Infusion pump flow rate (Flow<sub>infusion,i</sub>) (mL/h) | Amount infused (mg) |
| 1 | 17 | 7.1 | 0.446 | 1 | 25 | 7.1 | 0.621 | 1 | 24 | 5.0 | 0.575 |
| 2 | 19 | 6.5 | 0.446 | 2 | 29 | 6.3 | 0.621 | 2 | 27 | 4.4 | 0.575 |
| 3 | 20 | 5.9 | 0.446 | 3 | 33 | 5.4 | 0.621 | 3 | 32 | 3.8 | 0.575 |
| 4 | 23 | 5.3 | 0.446 | 4 | 39 | 4.6 | 0.621 | 4 | 38 | 3.2 | 0.575 |
| 5 | 26 | 4.7 | 0.446 | 5 | 48 | 3.7 | 0.621 | 5 | 47 | 2.6 | 0.575 |
| 6 | 29 | 4.1 | 0.446 | 6 | 63 | 2.9 | 0.621 | 6 | 61 | 2.0 | 0.575 |
| 7 | 34 | 3.5 | 0.446 | 7 | 89 | 2.0 | 0.621 | 7 | 88 | 1.4 | 0.575 |
| 8 | 42 | 2.9 | 0.446 | 8 | 154 | 1.2 | 0.621 | 8 | 162 | 0.7 | 0.575 |
| 9 | 52 | 2.3 | 0.446 |  |  |  |  |  |  |  |  |
| 10 | 71 | 1.7 | 0.446 |  |  |  |  |  |  |  |  |
| 11 | 112 | 1.1 | 0.446 |  |  |  |  |  |  |  |  |
| 12 | 276 | 0.4 | 0.446 |  |  |  |  |  |  |  |  |

\* ka and ke were empirically adjusted due to linezolid concentrations being too low after the first replicate

**Figure S3:**
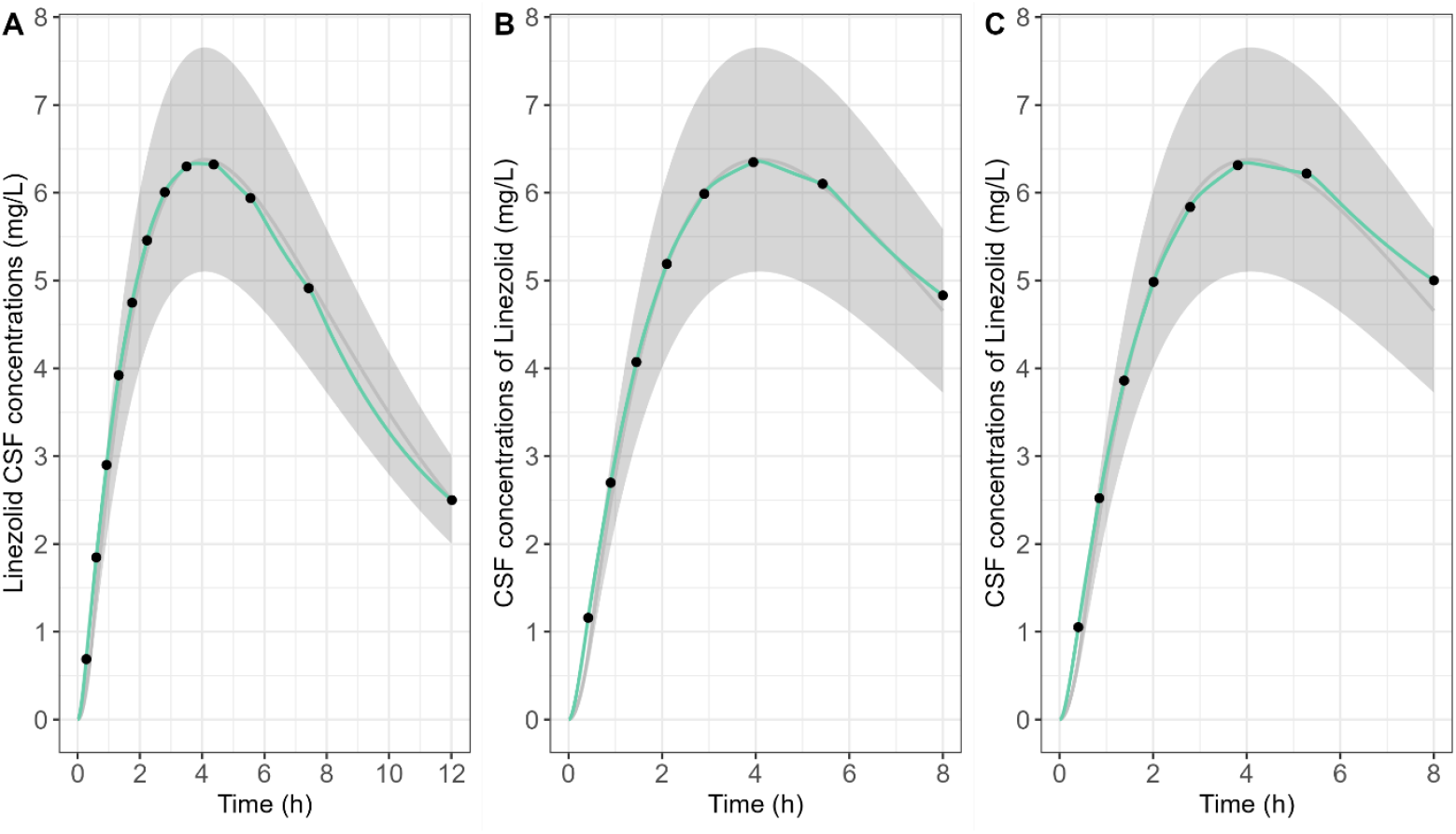
Target linezolid CSF concentrations from linezolid clinical population PK model (13) versus expected linezolid concentrations in the cartridge ECS for the first dose of 900 mg q12 h and 900 mg q8 h dosing regimen. (A) First dose of 900 mg q12 h. (B) First replicate of the first dose of 900 mg q8 h. (C) Second replicate of the first dose of 900 mg q8 h. The grey line corresponds to target linezolid concentrations. The grey area corresponds to a bias of 20 % from the target linezolid concentrations. The green line corresponds to expected linezolid concentrations in the cartridge ECS with the infusion pump program shown in **Table S3**. Black points correspond to the expected concentrations at the end of each infusion sub-interval.

